# Oligo DNA-based quantum dot (QD) single-particle tracking for multicolor single-molecule imaging

**DOI:** 10.1101/2025.10.06.680600

**Authors:** Shigeo Sakuragi, Naoki Kato, Tomoya Uchida, Boxiao Zhao, Taro Katagiri, Miyu Enomoto, Rie Kato, Hideaki Yoshimura, Chisato Oyama, Iona Katayama, Ayano Chikuma, Yuji Teramura, Hiroko Bannai

**Author notes:** Corresponding author: Hiroko Bannai, Waseda University, Shinjuku, Tokyo 162-0056, Japan.

## Abstract

Quantum dot single-particle tracking (QD-SPT), a powerful tool for analyzing membrane domains that are critical to various cellular processes, is widely used in membrane molecular dynamics research. QDs, which possess both a broad excitation range and a narrow emission bandwidth, are inherently suited for multicolor imaging at various wavelengths. However, applying multicolor QD-SPT with multiple biomolecular targets has been challenging due to the limited methods for specifically conjugating QDs to biomolecules. Here, we propose a DNA hybridization-based QD labeling method that generates several specific interactions based on sequence. QD fused with 20-mer oligo DNA specifically labeled membrane lipid 1,2-dipalmitoyl-sn-glycero-3-phosphatidylethanolamine (DPPE) covalently bound to complementary oligo DNA, forming a stable label that is suitable for SPT. The combination of polyA–polyT sequence as the linker oligo caused more QDs to fuse to DPPE compared with a random sequence linker. Oligo DNA-based QD-SPT accurately reflected the diffusion dynamics of DPPE measured using the single-fluorescence tracking technique and was compatible with conventional QD-SPT utilizing secondary antibody Fab. Using different pairs of oligo DNA sequences, we successfully achieved multicolor QD-SPT that distinguishes the lateral diffusion of DPPE and a membrane protein, GABA_A_ receptor (GABA_A_R), within the same cell. The oligo DNA-based QD labeling method developed in this study is anticipated to substantially advance simultaneous multicolor QD-SPT of different living cell membrane molecules specific to DNA sequences.

---

In the fluid-mosaic model proposed by Singer and Nicolson, biological membranes demonstrate fluid-like properties, in which membrane proteins and lipids that are embedded in the lipid bilayer undergo lateral diffusion across the plasma membrane [1]. However, high-resolution analyses at the sub-millisecond timescale have revealed that even lipid molecules that diffuse freely are transiently confined within compartments of approximately 13–100 nm in size, based on the actin cytoskeleton and transmembrane proteins anchored to it.

Lipid molecules display compartmentalized diffusion, characterized by restricted motion within individual compartments and occasional hopping between adjacent compartments [2] [3]. Furthermore, specific membrane proteins are localized in the membrane region on the nanometer to micrometer scale, generating functional membrane domains [4]. These domains are developed through fine regulation and restriction of lateral diffusion of membrane molecules and play crucial roles in a wide array of cellular processes, including signal transduction, synaptic transmission, and immune responses. For example, neurotransmitter receptors accumulate at the synapse, which serve as the fundamental units of inter-neuronal communication, and the number of receptors at the synapse, ultimately, the efficacy of synaptic transmission, is regulated via lateral diffusion constraints caused by interactions between receptors and scaffold proteins [5]. Further, transmembrane proteins and cytoskeletons restrict the lateral diffusion of membrane lipids in the axon initial segment of neurons, underlying the physical compartmentalization between neuronal input and output domains [6]. Observations of membrane molecule dynamics are crucial to understanding such self-organizing principles of plasma membranes and revealing the molecular mechanisms underlying their physiological functions.

Single-molecule imaging enables the observation of the dynamics of individual molecules, facilitating not only the assessment of ensemble behavior but also detailed information about the behavior of molecules in confined microdomains, stochastic movements, and heterogeneity within molecular populations. Single-particle tracking (SPT) is one of the representative methods in single-molecule imaging, in which probes are specifically conjugated to target molecules in live cells, causing the high-precision tracking of particle positions [7]. Among the SPT methods that use diverse probes, quantum dot (QD)-SPT has become a widely used approach for investigating the dynamics of membrane molecules in neurons. QDs are semiconductor nanocrystals that serve as fluorescent probes, which overcome limitations associated with organic fluorophores caused by their superior brightness and long fluorescence lifetimes [5]. Furthermore, their high brightness and significant Stokes shift provide an excellent signal-to-noise ratio (SNR), enabling more straightforward and higher-resolution analysis of fluorescence images compared with organic dyes [8]. However, QDs still have certain limitations, as the relatively large size of QD bioconjugates hinders access to spatially restricted regions, such as synaptic clefts, due to physical confinement [9] [10]. The blinking of QDs, i.e., intermittent fluorescence emission, hinders continuous tracking and quantitative analysis [11]. However, the optical advantages of QDs, particularly their high SNR, which helps improve spatial resolution, outweigh these drawbacks. Hence, QD-SPT has been extensively adopted in neuroscience research to investigate membrane molecule dynamics at the single-molecule level.

The most distinctive feature of QDs is that their fluorescence emission wavelength is precisely identified based on their particle size, leading to sharp and narrow emission spectra [12]. With their broad excitation range and narrow emission bandwidth, QDs are inherently well-suited for multicolor imaging of various wavelengths.

Indeed, studies that used fixed tissues and cells reported multicolor imaging employing QDs [13] [14] [15]. Despite developing QD-SPT over 20 years ago, reports that utilize QDs for multicolor imaging, particularly those involving labeling of multiple molecular species with QDs of distinct emission wavelengths, are limited. The lack of robust methods for specifically labeling different target molecules in living cells with QDs with different wavelengths of emission is one of the primary reasons. Currently, the commonly used QD labeling method employs the specific interaction between the primary antibodies and secondary antibodies or biotin–avidin interactions. However, these methods involve a small number of combinations of specific molecular interactions, which limits the options for multicolor imaging that simultaneously targets multiple different molecules. Therefore, novel QD-labeling strategies need to be developed to expand the applicability of multicolor QD-SPT in the same cell.

In recent years, multicolor immunofluorescence techniques employing deoxyribonucleic acid (DNA) barcodes have facilitated the visualization of multiple protein localizations within the same cell [16] [17]. In this study, we apply the specific and stable hybridization between complementary DNA strands to achieve persistent conjugation between membrane molecules and QDs. Unlike DNA-PAINT approaches that depend on rapid binding–unbinding kinetics [18], the present strategy is designed to provide stable probe attachment over the time scale of single-particle tracking. This enables sequence-specific targeting of distinct membrane molecules with QDs of defined emission properties. Utilizing this approach, we demonstrate a practical method for multicolor QD-SPT in the same cell, wherein multiple molecular species, the phospholipid and the neurotransmitter receptor, are labeled with QDs through 20-mer DNA hybridization, each emitting at different wavelengths.

## Materials and methods

### Primary culture of rat hippocampal neurons

In this study, all experiments were conducted following the guidelines approved by the Animal Experimentation Committee of Waseda University. Primary hippocampal neurons were prepared as previously described [19]. In brief, hippocampi were dissected from embryonic day 20 (E20) Wistar/ST rats (Japan SLC) and treated with 0.125% trypsin (Sigma-Aldrich) and 0.025 w/v% DNase (Sigma-Aldrich). The tissue was then dissociated in the plating medium, composed of the minimum essential medium (MEM; Thermo Fisher Scientific) supplemented with 2% MACS NeuroBrew-21 (Miltenyi Biotec), 2 mM L-glutamine (Nacalai Tesque), 1 mM sodium pyruvate (Nacalai Tesque), and 5 units/mL of penicillin and 5 µg/mL of streptomycin (Nacalai Tesque). Dissociated cells were seeded at a density of 1.4 × 10⁵ cells per well on an 18-mm polyethyleneimine-coated glass coverslip and cultured in a CO₂ incubator (37°C, 5% CO₂). Four days after plating, the culture medium was replaced with the maintenance medium, the Neurobasal-A (Thermo Fisher Scientific), supplemented with 2% MACS NeuroBrew-21, 2 mM L-glutamine, and penicillin (5 units/mL)/streptomycin (5 µg/mL). Cultures were maintained at 37°C in a CO₂ incubator until the experiments were performed at 14 days *in vitro* (DIV14) or later.

### Membrane molecule labeling using ssDNA

We prepared 20-base-pair ssDNA probes comprising one pair of complementary poly sequences (polyA and polyT) and two pairs of complementary random sequences (random1S and random1AS, random2S and random2AS). Supplementary Table S1 shows the sequence. For the QD-SPT of phospholipids with ssDNA-conjugated QDs, we prepared the 1,2-dipalmitoyl-sn-glycero-3-phosphatidylethanolamine (DPPE) tagged with ssDNA at the hydrophilic PEG end (ssDNA–DPPE) (Figure 1). As previously described, ssDNA was attached to maleimide-poly(ethylene glycol)-conjugated phospholipid DPPE through a thiol–maleimide reaction (Supplementary Figure S1A) [20]. In brief, DPPE and α-N-hydroxysuccinimidyl-ω-maleimidyl poly (ethylene glycol) (NHS-PEG-Mal, Mw: 5000) were combined to produce Mal-PEG-DPPE. Thiolated-labeled ssDNAs to the 5’ end were then reacted with Mal-PEG-DPPE to obtain the ssDNA–DPPE and maintained in the phosphate-buffered saline (PBS). Further, DPPE was conjugated with ssDNA via copper-free click chemistry through dibenzocyclooctyl (DBCO)–azide conjugation (ssDNA–DBCO–DPPE, Supplementary Figure S1B) [21] [22]. DPPE–PEG (2000)–azide (10 µM; Avanti) was reacted with 5’ DBCO-C6-N_2_-PolyA (20) (1 µM; Nihon Gene Research Laboratories) in PBS for 60 min at room temperature.

**Figure 1.**
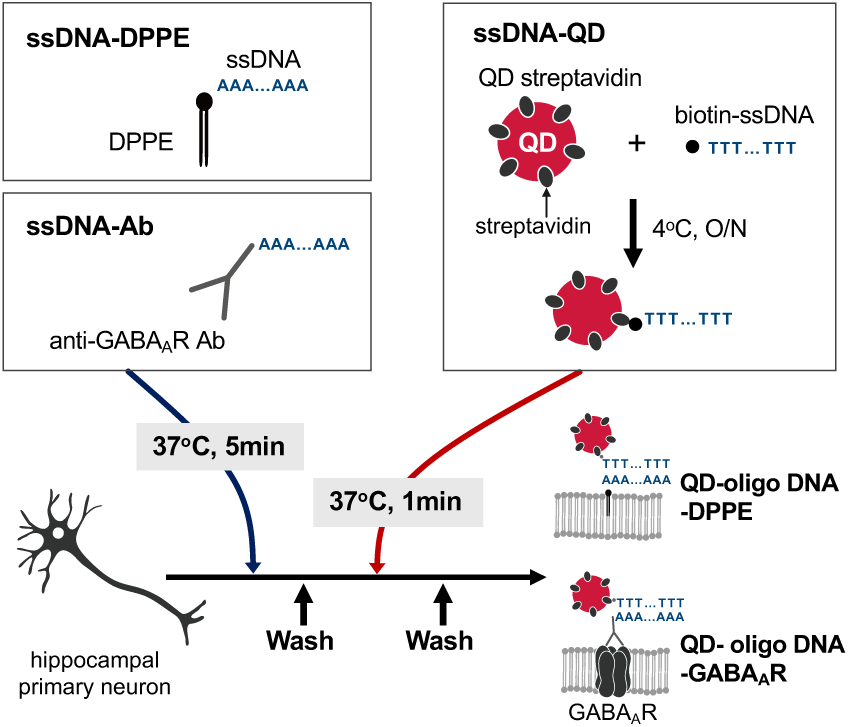
Overview of the ssDNA labeling protocol used in this experiment. DPPE conjugated with a 20-mer ssDNA (ssDNA–DPPE), or an anti-GABA_A_R antibody conjugated with DNA (ssDNA–Ab), and QDs modified with ssDNA via streptavidin–biotin interaction (ssDNA–QD) were prepared. Primary rat hippocampal neurons were incubated with ssDNA–DPPE or ssDNA–Ab for 5 min, followed by a 1-min incubation with ssDNA–QD, in a humid chamber maintained at 37°C.

Moreover, ssDNA and QD conjugates (ssDNA–QD) were prepared by mixing Qdot 655 streptavidin conjugate (1 µM; QD655; Thermo Fisher Scientific), ssDNA labeled with biotin at the 5’ end (10 µM), and Qdot Binding Buffer (QBB; 2.62 mM sodium tetraborate decahydrate [Wako], 39.5 mM boric acid [Nacalai Tesque], 2 w/v% bovine serum albumin [Sigma-Aldrich], 0.05 w/v% sodium azide) at a volume ratio of 1.0:0.5:8.5, followed by overnight incubation at 4°C. QD605 (Thermo Fisher Scientific) conjugated with ssDNA as described above was used for simultaneous labeling of multiple membrane proteins in parallel with ssDNA–QD655. The imaging medium, MEM without phenol red (Thermo Fisher Scientific), supplemented with 20 mM HEPES (Sigma Aldrich), 2 mM glutamine, 1 mM sodium pyruvate, and 33 mM glucose (Nacalai Tesque), was used for the dilution and the incubation of lipid and antibody, cell washing, and live-cell imaging. QBB was supplemented with 215 mM of sucrose (QBB-suc) to adjust the osmolarity when QDs were incubated with neurons.

To incorporate ssDNA–DPPE into the plasma membrane, ssDNA–DPPE (5 µg/mL) was incubated with primary cultured rat hippocampal neurons for 5 min. After washing, the coverslips were incubated with 0.063-0.25 nM ssDNA–QD in QBB-suc for 1 min, followed by washing and imaging in the imaging medium. Labeling processes, which include lipid, antibody, and QD incubation, were performed in a humid chamber placed on a heating plate maintained at 37°C [23].

For labeling GABA_A_ receptors (GABA_A_Rs) with ssDNA, we utilized a custom-made antibody that targets the extracellular domain of the γ2 subunit of GABA_A_R [24] conjugated with polyA (ssDNA–Ab; Figure 1). The antibody was conjugated to ssDNA at a 1:5 molar ratio employing the Oligonucleotide Conjugation Kit (Abcam, ab218260), which targets lysine residues via an amine-reactive ester. Endogenous GABA_A_R in cultured rat hippocampal neurons were labeled with ssDNA by incubating with 0.01 mg/mL of ssDNA–GABA_A_R antibody for 5 min, followed by incubation with 1 nM ssDNA–QD in QBB-suc for 1 min in a humid chamber.

### Conventional QD labeling using Fab fragment antibody

GABA_A_R labeling using Fab fragment antibody as a linker for QD (conventional labeling) was performed according to previous reports [25] [23]. Rat primary hippocampal neurons were incubated with a custom-made anti-GABA_A_R γ2 subunit antibody that targets the extracellular domain (0.01 mg/ml, 5 min) followed by incubation with biotinylated Fab fragment antibody (10 µg/ml, 5 min; Biotin-Fab, Jackson ImmunoResearch). After washing, GABA_A_Rs were labeled via biotin–avidin binding by incubation with 1 nM QD655 in QBB-suc at 37°C for 1 min.

### QD-SPT data acquisition

QD-SPT was conducted using an inverted microscope (IX73, Olympus) equipped with multi-wavelength LED illumination (niji, Bluebox Optics), an oil-immersion objective lens (×60, NA 1.42, Olympus), and filter sets for QD wavelength, i.e., an excitation filter (450/70 nm, e.g., Semrock, Rochester, NY; BrightLine® single-band band-pass filter #FF01-450/70-25), dichroic beam splitters (DM) and emission filter (EM) based on each QD emission wavelength (for Qdot655: 655/15 nm #FF01-655/15-25, DM #FF640-FDi01-25 × 36; for Qdot605: 605/15 nm #FF01-605/15-25, DM #FF593-Di03), and the EM-CCD camera (ImagEM C9100-13, Hamamatsu Photonics). MetaVue software (Molecular Devices) was used to control the excitation illumination and the EM-CCD camera. A total of 100 consecutive frames were acquired within a 41.8 × 41.8 µm field of view at 13 Hz with a 75 ms exposure time. During observation, the specimen and the microscope stage were maintained at 37°C using a heating system (Tokai Hit). In the DNase experiment, 0.05 w/v% DNase (Sigma-Aldrich) in Hank’s balanced salt solution (Nacalai Tesque) was added to the observation chamber on the microscope stage and left for 3 min. Images were then acquired in the same field of view after DNase administration.

### Single-fluorescence tracking for DPPE

The ATTO647–DPPE (ATTO-TEC GmbH, AD647-151) was inserted in the cultured neurons by incubating the coverslips with 0.1 µM ATTO647–DPPE diluted in the imaging medium for 10 min, within the humid chamber placed in a CO₂ incubator. After washing three times, the coverslip was mounted in the observation chamber and immersed in the imaging medium for observation.

Total internal reflection fluorescence microscopy (TIRFM) was used for single-molecule fluorescence imaging of organic dye ATTO647 [26], based on an inverted microscope (IX81, Olympus) equipped with 640-nm diode laser (CUBE, Coherent Inc.), 405/488/561/635-nm band excitation filter (Semrock Inc.), Dichroic mirror (FF662, Semrock Inc.), 697/58-nm band-pass filter (Semrock Inc.), EM-CCD camera (ImagEM, Hamamatsu Photonics), and an oil-immersion objective lens (×100, NA 1.49, Olympus). The imaging was performed in 256 × 256-pixel subarray mode, recording 200 frames at 30 Hz (33 ms/frame), at the surface and the bottom focal planes in the same field of view.

### Tracking analysis of QD and organic dyes

A tracking analysis of QD-SPT was conducted using TI Workbench as previously described [23] [25]. The center of each QD spot was defined based on cross-correlation of the fluorescent spots with a Gaussian model with a point spread function. QD-trajectories are automatically assembled by linking the centers of fluorescent spots. Only QDs with blinking were considered for the analysis to ensure that they are single particles, with the trajectories with > 30 points used for further analysis. The mean squared displacement (MSD) was calculated from the coordinates of reconstructed trajectories. The MSD-Δ𝑡 curve, which plots MSD versus time (Δ𝑡), was obtained using the following equation (1) [26]:

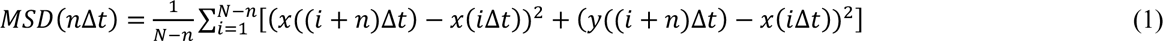

where 𝛥𝑡 denotes the acquisition time (75 ms), 𝑁 indicates the total number of frames, 𝑛 represents the frame number, and 𝑖 denotes the positive integers. The diffusion coefficient (D) of QDs was obtained by fitting the first 1–4 points without origin of the MSD-𝑛Δ𝑡 curves with the following equation (2):

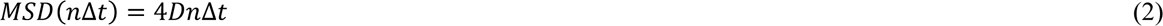

QDs with D of < 0.0004, equivalent to D of QDs adhered on the coverslips, were defined as “immobile” QDs [23].

Single-fluorophore tracking (SFT) images were filtered using the dilate (circle, 4 pixels) function in MetaMorph (Molecular Devices) to reduce spike-like noise, and the trajectories were composed employing TI Workbench software. To ensure single-molecule fluorescence, the analysis included only bright spots that disappeared in a single step during recording. The D values were calculated for trajectories tracked for more than 10 consecutive frames.

### Ca^2+^ imaging

Primary hippocampal neurons (DIV 14 or later), either with or without QD labeling for GABA_A_R and DPPE, were loaded with 0.5 µM Fluo-8 AM (AAT Bioquest) diluted with the imaging medium (MEM supplemented with 20 mM HEPES [pH 7.4], 2 mM glutamine, and 1 mM sodium pyruvate) for 5 min at 37°C. After washing, neurons were pre-incubated for 5 min in the imaging chamber with either muscimol (Sigma, 20 µM; GABA_A_R agonist) or imaging medium (mock control). The same conditions were maintained throughout the imaging session. Time-lapse imaging was performed at 1 Hz for 180 s using the same imaging configuration as for QD-SPT, with appropriate filter sets. Depolarization was induced by applying KCl (40 mM final concentration) 60 s after the start of recording, and imaging continued for an additional 120 s.

MetaMorph software (Molecular Devices) or ImageJ (NIH) was used for imaging analysis. Regions of interest (ROIs) were placed over 2–4 neuronal somata per experiment. Changes in intracellular Ca^2+^ levels were expressed as ΔF/F_0_, calculated as (F – F_0_)/F_0_, where F_0_ denotes the mean baseline fluorescence intensity before KCl application. The maximum ΔF/F_0_ after KCl stimulation was calculated for each ROI and compared between the muscimol-treated and mock-stimulated groups.

### Statistics

Complementary binding by the four ssDNA–QDs was compared using a one-way analysis of variance with a post hoc Tukey HSD test. The number of QDs before and after DNase application was compared using a Wilcoxon signed-rank test. A Kolmogorov–Smirnov test was used to perform the cumulative distribution of the D. Differences in D among three groups were analyzed using the Kruskal–Wallis test, followed by Dunn’s post hoc test. A Mann–Whitney *U*-test was used for other statistical analyses. The figure illustrates the *P*-values. Sample size (N) for each experiment is described in the figure legend. Each experiment utilized at least three independent culture batches.

## Result

### QD Labeling of Membrane Molecules depending on annealing between complementary ssDNAs

First, we investigated whether the phospholipid DPPE tagged with ssDNA (ssDNA–DPPEs) was specifically labeled with QDs conjugated with ssDNA (ssDNA–QDs) with a complementary sequence through the hybridization. The 20-mer oligo DNA, composed of identical bases or random sequences, was developed as ssDNA sequences (Supplementary Table S1) and added to QDs and target molecules. DPPEs conjugated with polyA ssDNA (polyA–DPPE) via the thiol–maleimide reaction (Supplementary Figure S1A) were inserted into the plasma membrane of primary rat hippocampal neurons and incubated with QDs conjugated with either polyA or polyT sequences (Figure 2A). QDs with polyA (polyA–QDs), carrying the same sequence as the ssDNA on the DPPE, did not remain on the surface of the neurons, whereas QDs with polyT (polyT–QDs), which carry the complementary sequence, were observed on the neurons (Figure 2B). The median number of QDs bound to neurons per image was 1 for polyA–QDs, whereas that for polyT–QDs was significantly higher at 241 per image (Figure 2C, Supplementary Table S2). Comparable to the result of polyA–DPPEs, the DPPE tagged with polyA by the copper-free click chemistry through the DBCO–azide reaction (polyA-DBCO-DPPE, Figure S1B) was targeted by polyT–QDs, but not by polyA–QDs (Supplementary Figures S1C, S1D), indicating that the linker between the DPPE and the ssDNA does not affect the DNA hybridization.

**Figure 2.**
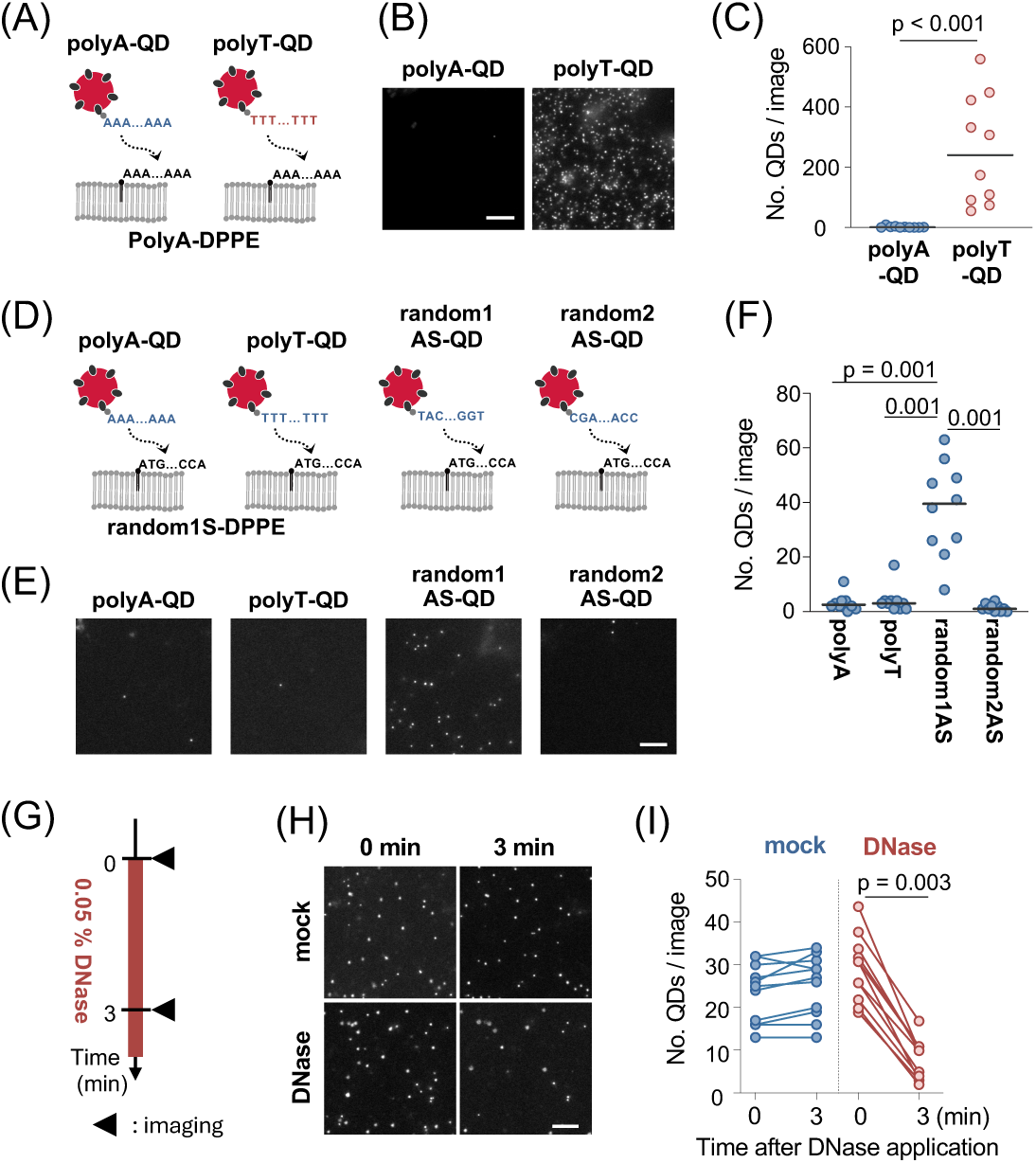
QD labeling of membrane lipids via complementary DNA sequences. (A) Schematic illustration of QD labeling of DPPE via complementary DNA hybridization. PolyA–DPPE was incorporated into the neuronal plasma membrane and incubated with either polyA–QDs or polyT–QDs. (B) Representative fluorescence images after labeling with polyA–QDs (left) or polyT–QDs (right). Scale bar indicates 10 µm. (C) Number of bound QDs per image. Solid lines indicate medians (N = 10 images per group). (D) Specificity test using random ssDNA sequences. Random1S–DPPE was probed with polyA–QD, polyT–QD, random1AS–QD (complementary), or random2AS–QD. (E) Representative images of the four ssDNA conditions. Scale bar indicates 10 µm. (F) Number of QDs per field of view. Solid lines indicate medians (N = 10 images per group). (G) DNase treatment time course. Images were acquired before and 3 min after adding 0.05% DNase or HBSS (mock). (H) Representative images of the same field before and after treatment. Scale bar indicates 10 µm. (I) Number of QDs per field of view (N = 11 images per group). Statistics: (C) Mann–Whitney U test; (F) one-way ANOVA with Tukey’s HSD; (I) Wilcoxon signed-rank test.

PolyA–QDs labeled the plasma membrane of neurons incubated with polyT–DPPE, whereas ssDNA–QD did not bind to neurons without polyT–DPPE (Supplementary Figures S2A, S2B). To evaluate nonspecific binding, neurons without polyT–DPPE were incubated with 0.063 nM polyA–QD or a 10-fold higher concentration (0.63 nM) (Supplementary Figure S2C). Fewer than one QD per field of view (41.8 × 41.8 µm) was detected at 0.063 nM, whereas the 10-fold higher concentration yielded 2–7 QDs per cell (4.4 ± 0.3, mean ± SEM) (Supplementary Figure S2D). Considering that more than 20 QDs (median) per field of view were typically detected under specific binding conditions (Supplementary Table S2), the number of non-specifically bound QDs remained substantially lower than the specific signal even at higher ssDNA–QD concentration. Furthermore, polyA–DPPE was rarely targeted with ssDNA–QD that carries two random sequences, random 1AS and random 2AS (Supplementary Figures S2E, S2F), indicating the high dependence of QD labeling via ssDNA on the DNA sequence complementarity.

To further confirm the binding specificity of QD labeling with ssDNA, we inserted a random ssDNA sequence-conjugated DPPE (random1S–DPPE) into the plasma membrane of cultured neurons and evaluated whether QDs with four different ssDNA sequences (polyA, polyT, random1AS, and random2AS) could bind to random1S–DPPE (Figure 2D). QDs with polyA, polyT, or random2AS, all of which are not complementary to random1S, did not remain on the cell surface. In contrast, a substantial amount of QDs with the complementary sequence random1AS bound to random1s–DPPE (Figure 2E, Supplementary Table S2). Statistical analysis revealed that random1AS–QDs were significantly more observed in the plasma membrane (Figure 2F,

Supplementary Table S2). These results indicate that hybridization between 20-base ssDNAs with complementary sequences exhibits sufficient binding specificity for QD labeling, even when using random sequences. Further, the binding between ssDNA–DPPEs and ssDNA–QDs was sensitive to DNase treatment (Figure 2G). Specifically, 0.05% DNase treatment of neurons labeled with random1S–DPPE and random1AS–QDs reduced the number of QDs in the same field, whereas the mock group remained unchanged before and after the addition of DNase (Figures 2H, 2I, Supplementary Table S2). This further emphasizes that the QD labeling was not nonspecific but instead oligo DNA-dependent. Collectively, the hybridization of ssDNAs with complementary sequences applies to the specific labeling of membrane molecules with QDs.

### Measurement of the diffusion coefficient (D) of DPPE by oligo DNA-dependent QD-SPT

The suitability of DPPE labeled with QDs was investigated through DNA hybridization for SPT. The diffusion of membrane molecules at the lower (coverslip-facing) membrane in some cell types is reduced compared with the upper (nonadherent) membrane [27] [28]. TIRFM was utilized to measure the diffusion coefficient (D) of the DPPE fused with the organic dye ATTO647 (ATTO647–DPPE) in the primary neurons at the upper and lower membranes of the same cell [26] (Figure 3A). ATTO647–DPPE trajectories at the upper and lower membranes demonstrated different diffusion properties (Figure 3B). The D of ATTO647–DPPE at the lower membrane (median: 1.15 × 10^−1^ µm²/s) and upper membrane (median: 2.21 × 10^−1^ µm²/s) were significantly larger than that of glass-attached ATTO647–DPPE (median: 2.40 × 10^−2^ µm²/s, Figure 3C and Supplementary Table S2). Furthermore, the D at the lower membrane was significantly lower than that at the upper membrane (Figures 3C and 3D, Supplementary Table S2), indicating the reduced lateral mobility of DPPE at the lower membrane of the primary cultured neuron.

**Figure 3.**
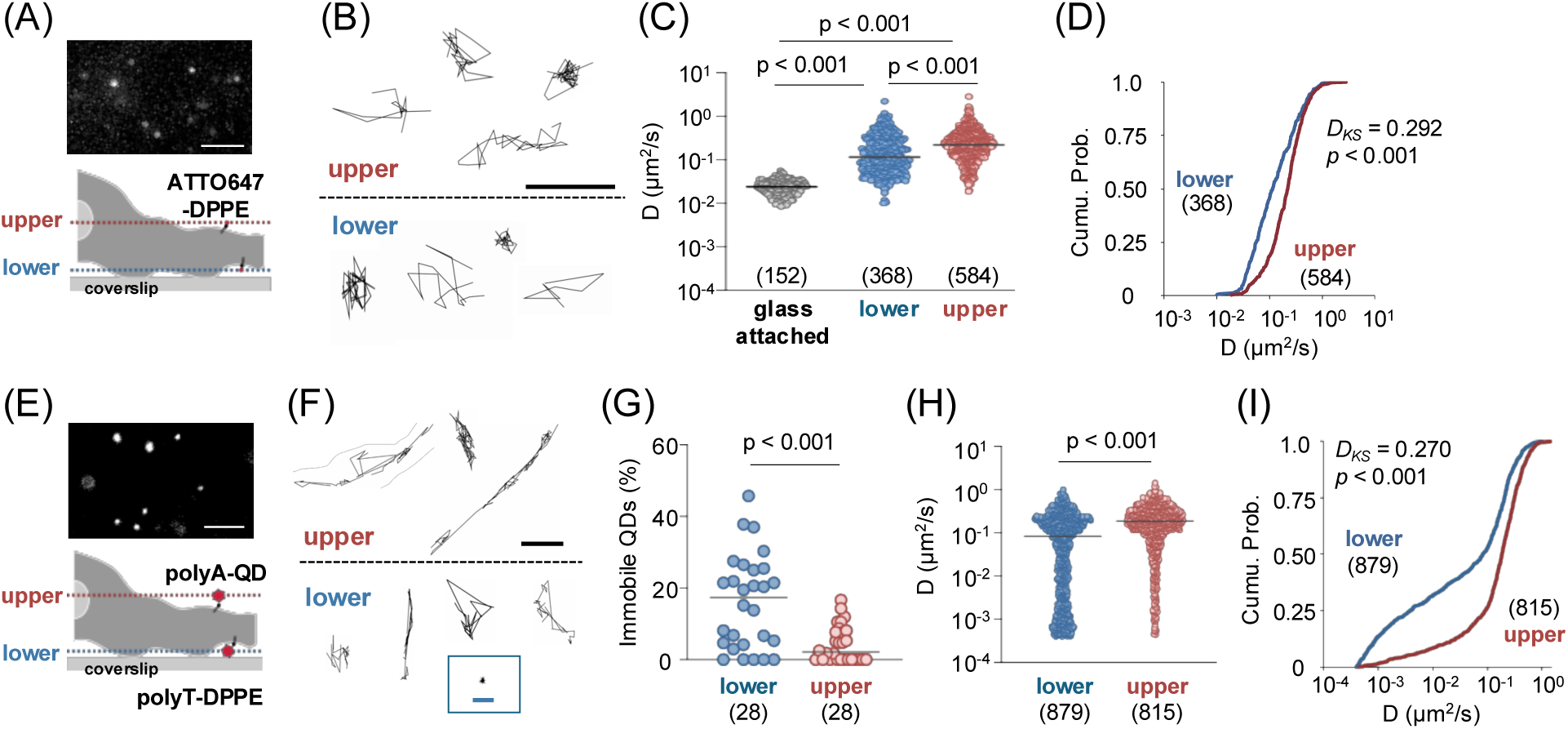
Diffusion dynamics of the DPPE measured using single-fluorophore imaging and oligo DNA-based QD-SPT. (A) ATTO647–DPPE labeling scheme and representative upper-membrane image. Scale bar, 5 µm. (B) Representative trajectories of ATTO647–DPPE at the upper and lower membranes. Scale bar, 1 µm. (C) D of ATTO647–DPPE at the glass-attached (gray), lower (blue), and upper (red) membranes. (D) Cumulative distributions of D at the upper and lower membranes. (E) OligoDNA-based QD labeling scheme and representative upper-membrane image. Scal bar indicates 5 µm. (F) Representative QD trajectories at the upper and lower membranes. Scale bars indicate 1 µm (main) and 0.2 µm (inset). (G) Fraction of immobile QDs per field of view. (H, I) Dot plot (H) and cumulative distribution (I) of D of mobile QDs. Statistical analysis: (C) Kruskal–Wallis test with Dunn’s post hoc test; (G, H) Mann–Whitney U test (Solid lines indicate medians); (D, I) Kolmogorov–Smirnov test (*Dks,* KS distance). N is reported in parentheses as spots (C, D), images (G), and QDs (H, I) (see Supplementary Table S2).

The diffusion of polyT–DPPE labeled with polyA–QDs was then measured at the upper and lower membranes of cultured neurons (Figure 3E). QD–polyA/polyT–DPPE complex trajectories exhibited lateral diffusion on the plasma membrane (Figure 3F), indicating the sufficient stability of QD labeling via oligo DNA for tracking membrane diffusion dynamics. The percentage of immobile QDs, QDs with D of less than 4.00 × 10^−4^ µm²/s [23], was significantly larger at the lower membrane (median: 17.3%) than at the upper membrane (median: 2.16%) (Figure 3G, Supplementary Table S2). Furthermore, the D of mobile QD–ssDNA–DPPE remained smaller at the lower membrane (median: 0.82 × 10^−1^ µm²/s) than at the upper membrane (median: 1.86 × 10^−1^ µm²/s), with a greater proportion of low D at the lower membrane (Figures 3H and 3I, Supplementary Table S2). These results indicate a suppressed diffusion of the QD–oligo DNA–DPPE complex at the lower membrane of cultured neurons, in coordination with the finding of ATTO647-conjugated DPPE whose molecular size is smaller than that of QDs (Figures 3C and 3D). However, a more immobile QD fraction and a smaller D at the lower membrane indicate the existence of steric hindrance due to the QD’s relatively large size compared with organic dyes. These results indicate that molecules in the upper membrane layer, where the effect of physical diffusion suppression is minimal, should be considered in subsequent experiments measuring D.

### Impacts of ssDNA Sequence on QD labeling efficacy and diffusion coefficient (D)

We examined whether ssDNA sequence affects the QD binding affinity and D with the ssDNA sequence pair of polyA and polyT (melting temperature [Tm] 52.17°C) and random1S/1AS (Tm: 64.47°C) (Figure 4A). The same ssDNA–DPPE (5 µg/mL) and ssDNA–QD (0.25 nM) concentration resulted in a higher number of QDs per image in the polyA/polyT ssDNA pair group than the random1S/1AS pair (Figure 4B, Supplementary Table S2). Despite having the lower Tm, polyA/polyT combination detected ∼10 fold more QDs (median:193 QDs/image), compared with random1S/1AS (median: 20 QDs/image) (Figure 4C, Supplementary Table S2). This indicates that the poly ssDNA sequence has higher labeling efficacy compared with the random sequence under current experimental conditions. QD-SPT should be performed at a sufficiently low density so that each QD can be identified as a single particle optically without overlapping the trajectory [9]. Therefore, ssDNA–DPPE and ssDNA–QD concentrations were adjusted based on the oligo DNA pair sequence in experiments conducted thereafter; thus, fewer than ∼50 QDs are found in one field of view (polyA/polyT pair: 1 µg/mL ssDNA–DPPE and 0.063 nM ssDNA–QD; random sequence pair: 20 µg/mL ssDNA–DPPE and 0.25 nM ssDNA–QD).

**Figure 4.**
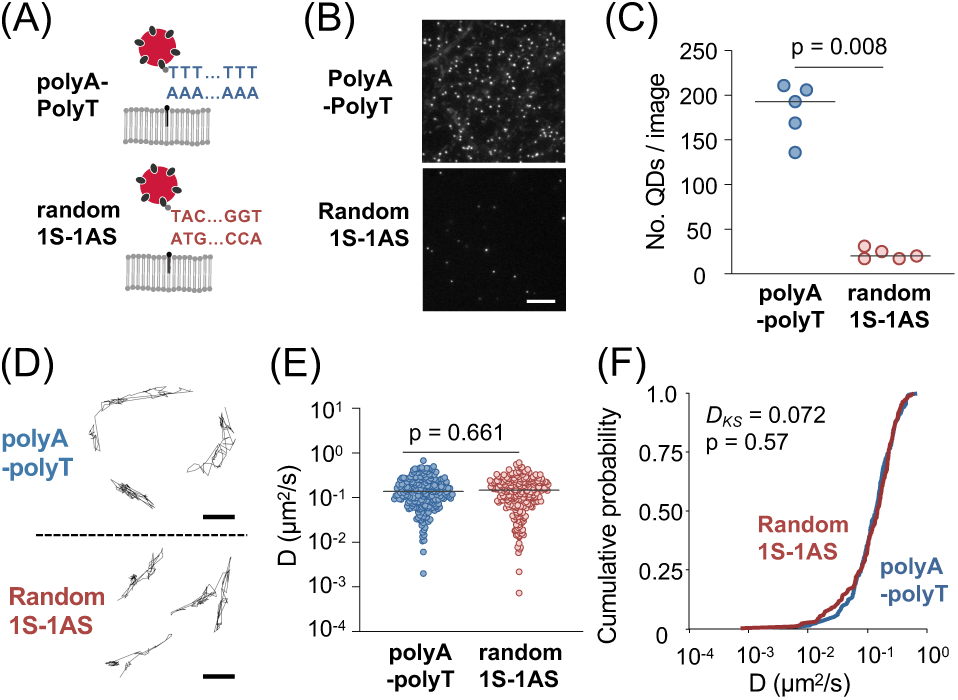
Impact of oligo DNA sequence on the diffusion dynamics of QD–oligo DNA–DPPE. (A) QD labeling via complementary sequences of polyA/polyT or random ssDNA. (B) Fluorescence images of neuron surface labeled with 0.25 nM polyT–QD and 5 µg/mL of polyA–DPPE (top) or 0.25 nM random1AS–QD and 5 µg/mL of random1S–DPPE (bottom). Scale bar indicates 10 µm. (C) Number of QDs per field. The line indicates the median. N = 5 images for both groups. (D) Example of trajectories of QD labeled via polyA/polyT (top) or random ssDNA (bottom). Scale bar indicates 1 μm. (E, F) The dot plot (E) and cumulative probability distribution (F) of D for QDs labeled with polyA/polyT or random sequence oligo DNA. Solid lines on the plot indicate medians; N = 220 (poly) and 257 (random) QDs. Mann–Whitney U-test for (C and E) and Kolmogorov–Smirnov test for (F).

Under our experimental conditions, each QD is expected to carry more than one ssDNA. To investigate whether such structural multivalency might promote clustering of ssDNA-DPPE on the plasma membrane, we conducted a quantitative binding analysis for the random1S/1AS pair and an independent random sequence (random2S/2AS) (Supplementary Figure S3). In this assay, the QD–ssDNA concentration was held constant at 0.25 nM, whereas the amount of ssDNA-DPPE applied to cells varied over a wide range (0–100 µg/mL). Binding increased with ligand concentration and reached a clear plateau (Bmax = 208.6 for random1S/1AS; 224.8 for random2S/2AS, Supplementary Figures S3B and S3D), consistent with a finite number of accessible membrane binding sites. Under the experimental conditions utilized throughout this study, the observed number of bound QDs per cell (∼50) was well below this saturation level (<25% of Bmax). Further, the working concentration of ssDNA-DPPE (20 µg/mL) remained below the half-maximal binding concentration (C_1/2_ = 24.73 µg/mL for random1S/1AS; 27.98 µg/mL for random2S/2AS, Supplementary Figures S3B and S3D), indicating that membrane occupancy was maintained in a sub-saturating regime. Under these sparse labeling conditions, the likelihood that a single QD bridges multiple spatially separated ssDNA-DPPE molecules is expected to be low. We cannot formally exclude rare multivalent interactions at the single-particle level; however, the quantitative binding behavior argues against substantial QD-induced aggregation under our imaging conditions.

To investigate the association of ssDNA sequences with the diffusion dynamics of target molecules, we compared the diffusion dynamics of DPPE labeled with QDs conjugated to either polyA/polyT pair or random1S/1AS pair. The trajectories of QD–oligo DNA–DPPE complex of both polyA/polyT pair and random1S–AS pair exhibited similar diffusion behavior on the plasma membrane (Figure 4D). No significant difference in the median and distribution of D was found between the polyA/polyT pair and the random1S/1AS pair (Figures 4E, 4F, Supplementary Table S2). These results indicate that the sequence of linker ssDNA affects labeling efficacy but has little effect on the D of oligo DNA-based QD-SPT.

### DNA-based QD-SPT for membrane proteins comparable to conventional QD-SPT

We further applied oligoDNA-dependent QD labeling to the diffusion analysis of a membrane protein. Here, for the proof of concept, we selected an inhibitory neurotransmitter receptor GABA_A_R, whose diffusion behavior on the neuron has been analyzed using conventional QD-SPT [25] [29] [30] [31]. PolyA ssDNA was conjugated to the anti-GABA_A_R antibody (polyA-Ab) and targeted to the endogenous GABA_A_Rs in the neuronal plasma membrane, followed by the incubation with polyA–QD or polyT–QD (Figure 5A). Only polyT–QD, which are complementary to the polyA sequence, bound to polyA–GABA_A_R (Figure 5B). The median number of QDs per image was 2 for polyA–QD and 28 for polyT–QD (Figure 5C, Supplementary Table S2). These results indicate that QDs can be specifically targeted to the protein of interest via oligo DNA hybridization.

**Figure 5.**
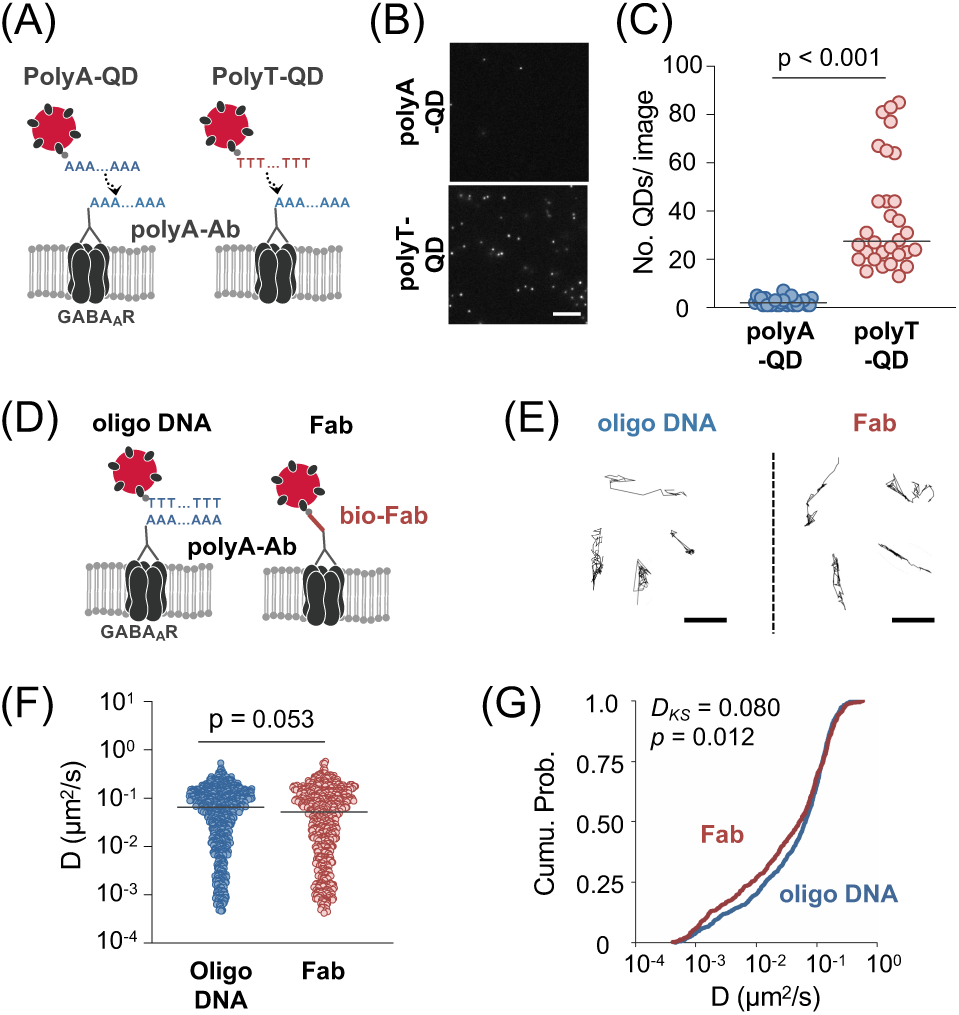
Analysis of membrane proteins with oligo DNA-based QD-SPT and conventional QD-SPT. (A) Labeling of QD with antibodies against GABAAR γ2 subunit conjugated with polyA ssDNA (polyA–Ab). PolyT–QD and polyA–QD were targeted to the polyA–Ab. (B) Representative QD images on neurons with polyA–Ab, after incubating with polyA–QD (top) or polyT–QD (bottom). Scale bar indicates 10 µm. (C) Number of QDs per field. Solid lines indicate the median. n = 30 images for both groups. (D) Schematic diagram of GABA_A_R QD labeling with oligo DNA-based labeling (oligo DNA) and conventional labeling with secondary antibody Fab fragments (Fab). (E) QD trajectories for oligo DNA-based labeling (left) and the conventional labeling (right). Scale bar indicates 1 µm. (F, G) Dot plot with the median solid lines (E) and cumulative probability distribution (F) of D for oligo-DNA-based or conventional DNA labeling. N = 969 (oligo DNA) and 686 (Fab). Mann–Whitney U-test for (C and F) and Kolmogorov–Smirnov test (G).

To investigate whether QD labeling affects GABA_A_R function at the cellular level, we conducted Ca^2+^ imaging experiments under muscimol stimulation. The application of the GABA_A_R agonist muscimol in unlabeled neurons significantly suppressed the increase in intracellular Ca²⁺ induced by 40 mM KCl application that induced the activation of voltage-dependent Ca^2+^ channels, consistent with GABA_A_R-mediated hyperpolarization (Supplementary Figure S4A). Neurons labeled with QD-ssDNA-GABA_A_Rs (Supplementary Figure S4B), as well as cells with QD-ssDNA-DPPE (Supplementary Figure S4C), demonstrated a similar suppression of KCl-induced Ca^2+^ elevation in the presence of muscimol treatment (Supplementary Figure S4D). These results indicate that GABA_A_R-mediated inhibitory signaling is preserved at the cellular level after QD labeling.

We then compared oligo DNA-based QD-SPT with conventional QD-SPT, which targets QDs via secondary antibody Fab. PolyA-Abs on the GABA_A_R were labeled with polyT–QDs (oligo DNA) or with a conventional method using biotinylated Fab antibodies and streptavidin–QDs (Fab) (Figure 5D). In both methods, QD-labeled GABA_A_Rs demonstrated diffusive behavior on the neuronal membrane (Figure 5E), with no statistically significant difference in the median D (Figure 5F, oligo DNA: 6.49 × 10⁻² µm²/s; Fab: 5.19 × 10⁻² µm²/s).

However, analysis of the distribution of D revealed a tendency for D-values to be high in the slow fraction of QDs–oligo DNA–GABA_A_R (Figure 5G; Kolmogorov–Smirnov test, *p* = 0.012, *D_KS_* = 0.080). Collectively, oligo DNA-based labeling functions as an effective alternative method to secondary antibody Fab labeling in QD-SPT of membrane proteins. However, differences in the linkers between QD and protein may affect the distribution of D.

### Multicolor imaging of the dynamics of lipids and proteins with oligo DNA-based QD-SPT

The results obtained so far demonstrate that oligo DNA-based QD-SPT can be applied to both membrane lipids and membrane proteins, yielding results equivalent to those obtained using conventional labeling methods. Finally, we investigated whether oligonucleotide-based QD-SPT enables simultaneous measurement of the diffusion of multiple membrane molecules within the same neuron using spectrally distinct QDs.

Before labeling cells, we verified that QDs of different wavelengths could be reliably distinguished under our imaging conditions. The QD605 and QD655 probes were mixed, immobilized on glass, and imaged within the same field of view using the same acquisition settings applied to both probes. Under these conditions, QD605-labeled particles were not detectably observed in the QD655 detection channel. Conversely, QD655 signals were not detected under QD605 imaging conditions (Figure 6A). This indicates negligible channel cross-talk in our dual-color imaging setup (Figure 6A).

**Figure 6.**
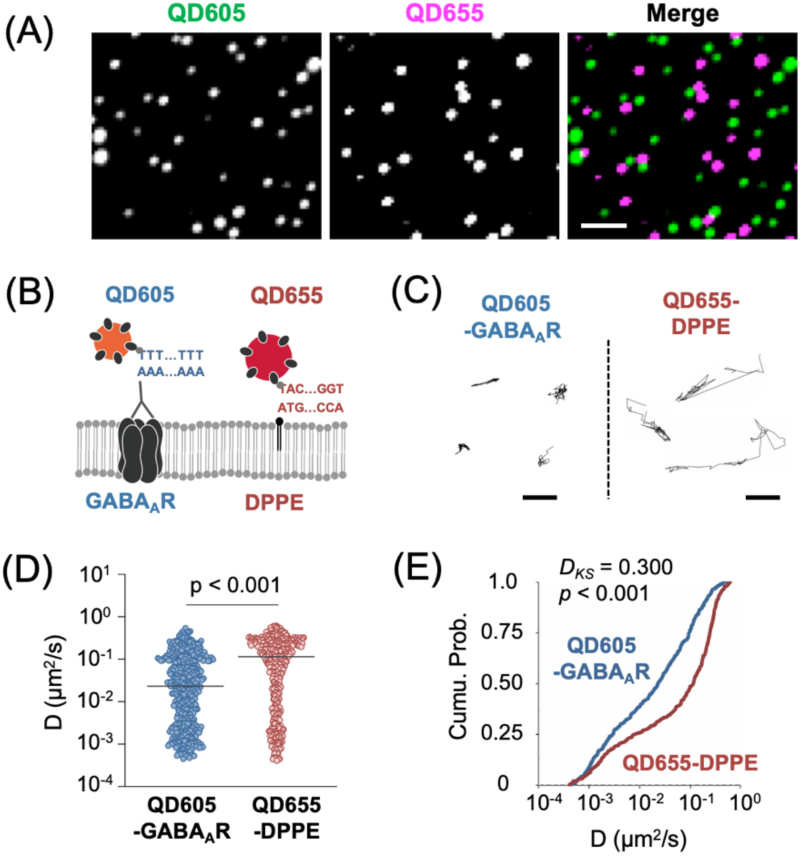
DNA-mediated two-color QD-SPT of and DPPE. (A) Images of QD605 and QD655 attached to glass, detected using wavelength-specific filters. (B) Strategy for multicolor QD labeling of GABA_A_R with QD605 and DPPE with QD655 using oligoDNAs. (C) Representative trajectories of QD605–GABA_A_R (left) and QD655–DPPE (right). Bars, 1 µm. (D, E) Distribution of D for QD605–GABA_A_R and QD655–DPPE shown as a dot plot (solid lines, medians) (D) and cumulative distribution (E). n = 419 (QD605–GABA_A_R) and 412 (QD655–DPPE). Statistical analysis: (D) Mann–Whitney U test; (E) Kolmogorov–Smirnov test.

GABA_A_R were then targeted with polyA–Ab and polyT–QD605 (GABA_A_R–QD605), and random1S–DPPE were labeled with random1AS–QD655 (DPPE–QD655) (Figure 6B). To assess potential cross-effects between QD labeling of different molecular species, we compared single- and double-label conditions using neurons from the same batch (Supplementary Figure S5A and S5B). The trajectories and D distributions for DPPE under double-label conditions (DPPE + GABA_A_R) were indistinguishable from those in single-label experiments (Supplementary Figure S5C–S5E). Similarly, GABA_A_R trajectories and D distributions were unchanged under double-label conditions (Supplementary Figure S5F–S5H). No statistically significant differences were observed between single- and double-label conditions for either molecule (Mann–Whitney U test and Kolmogorov–Smirnov test; n ≥ 298 trajectories per condition; *p* > 0.05). These results indicate that labeling one molecular species does not measurably change the diffusion dynamics of the other within the same cellular context.

Finally, trajectories of GABA_A_R–QD605 and DPPE–QD655 were analyzed. Representative 100-frame trajectories (∼8 s) of GABA_A_R–QD605 appeared shorter than those of DPPE–QD655 (Figure 6C), suggesting different diffusion behaviors. DPPE–QD655 (median: 1.15 × 10⁻¹ µm²/s) exhibited a significantly higher D than GABA_A_R–QD605 (median: 2.32 × 10⁻² µm²/s) (Figure 6D and Supplementary Table S2), and their D distributions were clearly distinct (Figure 6E; Kolmogorov–Smirnov test, *p* < 0.001, *Dks* = 0.300). The median D and interquartile ranges obtained using single-color QD-SPT were comparable to those obtained with multicolor QD-SPT (Supplementary Table S2). These results demonstrate that two-color QD-SPT using ssDNA enables simultaneous targeting and discrimination of two distinct membrane molecules based on QD emission wavelength.

## Discussion

In the present study, we demonstrated a method for labeling membrane molecules with QDs using the hybridization between ssDNAs with complementary sequences. This oligo DNA-based QD labeling was stable enough for tracking the dynamics of membrane molecules and compatible with conventional QD-SPT using a secondary antibody Fab. Furthermore, using different ssDNA sequences to label multiple types of membrane proteins with QDs of various wavelengths, we established multicolor QD-SPT, which enables observation of the dynamics of multiple membrane molecules, i.e., DPPE and GABA_A_Rs, within the same cell.

### QD labeling of membrane molecules via oligo DNA and dependence on the sequence

QDs exhibit a broad absorption spectrum and a narrow fluorescence spectrum, enabling excitation with a single light source and separation of multiple fluorescence signals. These QD characteristics are desirable properties for multicolor imaging, which enables the observation of different targets at various wavelengths in the same cell. Studies have labeled multiple types of molecules with QDs of different wavelengths in fixed cells [13] [14] [15]; however, most targets of QD-SPT in live cells were a single type of molecule. One of the reasons for this limitation was the combination of linkers connecting the membrane molecules and QD, such as the animal species of antibodies, and the number of protein-ligand interactions. In this study, we aimed to overcome this limitation of the number of combinations by applying hybridization between ssDNA, which generates numerous specific interactions based on the sequence. All 20 base oligo DNAs designed in this study, polyA/polyT pair, random1S/1AS pair, and random2S/2AS pairs, provided sufficient binding affinity and specificity, enabling hybridization-dependent labeling with QDs to DPPE molecules inserted in the plasma membrane. However, the number of QDs bound differed according to the sequence of linker oligo DNA. More QD bindings were observed when poly ssDNA was used as a linker, compared with random sequence. Under our experimental condition at 37°C, over 10-fold more QDs were found with polyA/polyT pair without guanine–cytosine pair than with random1S/1AS pair which has a higher Tm, although lower DNA duplex stability was expected for polyA/polyT combination (Tm: 52.17°C) compared with random1S/1AS pair (Tm: 64.47°C) [32]. One explanation for the higher binding of QDs by polyA/polyT pair is the high degree of freedom in sequence overlap, enabling stable binding even if only part of the 20-mer hybridizes. Further, ssDNAs with a random sequence can form secondary structures through base pairing. Indeed, secondary structure prediction of random ssDNA sequences using mFold [33] indicated the formation of secondary structures, which may inhibit the access to the complementary strand (Supplementary Figure S6), resulting in fewer observed QD counts. The minimum free energy change (*ΔG*) for these secondary structure formations is −1.74 kcal/mol or higher. Conversely, the free energy change during hybridization of complementary strands of random sequences calculated using DINAmelt [34] was −20 kcal/mol or higher, indicating that hybridization between complementary strands is more stable compared with secondary structure within the ssDNA. The DNA hybridization time was set to 1 min in the present study; however, extending the incubation time to reach equilibrium could potentially yield different results for the observed QD counts. However, the diffusion parameters, such as *D*, were not affected by the ssDNA sequence under conditions where the number of QD bindings per field of view was adjusted to be suitable for SPT. These results indicate that both poly ssDNA and random ssDNA are available for oligo DNA-based QD-SPT linkers.

Under the experimental conditions described here, the most crucial parameter for achieving an optimal QD labeling density was the surface density of ssDNA moieties that are directly associated with the plasma membrane (e.g., ssDNA-DPPE or ssDNA-conjugated anti-GABA_A_R antibody). When the ssDNA-QD concentration is kept constant, and the number of detected particles approaches the binding plateau, reducing the membrane-associated ssDNA density is an effective first step to prevent over-labeling and minimize potential multivalent crosslinking. Further, appropriate sequence design can tune hybridization efficiency, as random sequences typically demonstrate lower binding efficiency than polyA–polyT pairs. Thus, selecting the ssDNA sequence provides an additional means to modulate labeling density. Further reduction of ssDNA-QD concentration can help fine-tune particle density for single-particle tracking if membrane ssDNA density remains high after these adjustments.

### Pros and cons of oligo DNA-based QD-SPT against SFT

As reported in other cultured cells [27] [28], the diffusion dynamics of phospholipids vary between the surface and bottom layers of primary cultured rat hippocampal neurons. The *D* of DPPE labeled with ATTO647 was lower in the bottom layer than in the surface layer. Further, the *D* at the cell bottom obtained by oligo DNA-based QD-SPT was significantly lower than that in the surface layer, with an increased proportion of immobile QDs. Although limited to mobile QD fractions, the *D* of DPPE measured in the bottom layer was significantly lower than that in the surface layer. These results indicate that oligo DNA-based QD-SPT can reliably detect differences in diffusion dynamics at the subcellular level. However, only QD–oligo DNA–DPPE exhibited lower *D* (< 10⁻³ µm²/s) and immobile fractions (D < 0.0004 µm²/s) in the bottom layer, which were not observed in the ATTO647–DPPE group. The size of organic dyes is approximately 1–2 nm, whereas QDs with functionalization layers are approximately 15–30 nm [35]. Therefore, interactions between the QD as a probe and glycans or the extracellular matrix may affect the lateral diffusion of cell membrane molecules. The diffusion of QD-conjugated membrane proteins, calcium-activated potassium channels and B cell receptors, is significantly reduced at the bottom layer of the cell compared with proteins labeled with organic dyes [36] [37]. The difference in diffusion motion observed in this study between ATTO647-labeled DPPE and QD-labeled DPPE at the bottom layer of the cell may indicate the existence of steric hindrance that affects phospholipid diffusion. However, notably, the localization accuracy for single molecules decreases in single-fluorescence tracking due to the low photon count[38]. Indeed, the apparent D of ATTO647-DPPE immobilized on glass was 2.4 × 10^-2^ µm^2^/s. These molecules are physically immobilized; thus, this nonzero value reflects the localization precision and analytical limit of the single-fluorophore tracking method. These results indicate that D at or below this value cannot be reliably distinguished from measurement noise in the SFT approach. The main advantages of QDs are their bright fluorescence, which provides a high SNR, and their long fluorescence lifetime, which enables stable tracking. QD-SPT is a practical choice in environments where small spatial obstacles are expected, and when high localization accuracy and long trajectories are required.

### Effect of linkers on the protein diffusion measured by QD-SPT

We used the GABA_A_R for the example of the membrane proteins to compare oligo DNA-based QD-SPT with the widely used conventional QD-SPT using Fab fragments. The QD labeling achieved through ssDNA hybridization also maintained sufficient specificity and stability for SPT, such as labeling using the interaction between primary and secondary antibodies. The *D* of GABA_A_R showed no significant difference in their median values between the oligo DNA-based QD-SPT and the conventional QD-SPT. However, in the oligo DNA-QD-SPT method, the proportion of QDs showed significantly higher *D* in the fraction with low D values (K–S test, *p* = 0.012) compared with QD conjugated through Fab. The impact on the overall distribution of *D* was small (K–S test *Dks* value = 0.080); however, the potential influence of the linker between QDs and target molecules should be fully considered when combining the DNA method and the antibody approach for membrane proteins.

The labeling using annealing between complementary ssDNA is an effective method for increasing the number of specific intermolecular bonds; however, the process of fusing oligoDNAs to primary antibodies has been a major challenge. Advancements in copper-free click chemistry, which was utilized for the DPPE label in this study, have profoundly transformed the landscape of the field of bioconjugation. The increased availability of modified oligoDNAs and reagents for click chemistry enabled the fusion of oligonucleotides to antibodies and biomolecules in a few steps at room temperature via chemical reactions in biological laboratories [21]. Therefore, protein labeling using oligonucleotides as linkers is expected to be an option in the future.

The hydrodynamic radius of the Fab antibody is approximately 2.5 nm [39], and the hydrodynamic radius of the 20-base double-strand DNA is estimated to be 4–6 nm [40], indicating the comparability of the overall size of the Fab-QD and DNA-QD probes. The root mean square deviation calculated by molecular dynamics simulations is of the same order (1–5 Å), [41] [42] [43] [44], suggesting that differences in the size or rigidity of the linker molecules cannot explain the difference in diffusion dynamics. The mechanism by which differences in the linker result in variations in the D remains to be clarified in future studies.

### Prospects for Multicolor QD-SPT

QDs with sharp fluorescent spectra are ideal probes for multicolor imaging. Labeling the same membrane molecules with QDs of eight wavelengths helps avoid trajectory overlap in the same field of view, enabling tracking of the dynamics of more QDs, thereby significantly increasing the throughput of QD-SPT [45]. In the present study, we aimed to develop a new labeling method that utilizes the sequence specificity of ssDNA hybridization to target QDs of different wavelengths to specific targets. We successfully fused QD655 and QD605 to different targets, namely lipids (DPPE) and membrane proteins (GABA_A_R), respectively. The biotin–streptavidin interaction used for linking QD and oligoDNA forms a firm non-covalent bond (dissociation constant Kd =10^−15^ M) [46], and the possibility of dissociation is expected to be extremely low. The biotin–streptavidin binding was sufficiently stable not to allow cross-talk between QDs, considering the finding that DPPE and GABA_A_R diffusion, both of which have significantly different D values, could be distinguished using QD655 and QD605.

Here, we performed a two-color QD-SPT as a proof of concept for the availability of oligo DNA for live-cell multicolor imaging. Previous studies using fixed cells or fixed tissues have reported the application of QDs to multicolor detection methods. Flow cytometry was performed using simultaneous detection of 17 fluorescent probes, utilizing seven QDs and ten organic dyes [47]. Further, in single-cell imaging, a study revealed the staining of human prostate cancer cells using five types of biomolecules with five different QDs (QD525, QD565, QD605, QD655, and QD705) [48] [49]. This method requires dividing the labeling process into multiple steps and repeating the blocking of the antigenic sites of the primary antibody with a secondary antibody at saturation levels as many times as required. This approach is not ideal for SPT of live cells, which involves labeling to be performed quickly and at low density. In contrast, oligo DNA-based QD-SPT uses oligo DNA as linkers, which generates several combinations due to their sequence diversity, thereby enabling simultaneous labeling of multiple target molecules with different-colored QDs in a short time (several minutes). In principle, multicolor QD-SPT with three or more colors in the same cell is considered feasible; however, experimental verification remains a future challenge.

To perform multicolor QD-SPT, the optical system needs to be optimized using the sharp spectrum of QDs. In this study, the same field of view was recorded separately at each wavelength. Still, an optical system that simultaneously records multiple colors of QDs is desirable to be developed to minimize observation time. An image-splitting optical system can be employed to split the emission into two wavelengths and focus them onto a single camera, thereby enabling simultaneous imaging of two colors [50]. Furthermore, using two cameras and an image-splitting optical system, four colors of QDs within the same field is possible to be simultaneously imaged.

## Conclusion

In this study, we revealed that a linker using hybridization between complementary ssDNA can be utilized for the analysis of membrane molecular dynamics with QD-SPT. The DNA-based QD labeling method developed in this study is expected to help target multiple types of molecules with different wavelength QDs in an oligo DNA sequence-specific manner. It is considered to significantly enable simultaneous multicolor QD-SPT of living cell membrane molecules.

## Author contributions

SS, TK, NK, BZ, TU, ME, RK, HY, HB conducted the experiment. SS, TK, NK, TU, BZ, ME, RK, CO, IK, and AC performed the analysis. YT synthesized the ssDNA–DPPE. HB designed the study and supervision of this work. All authors discussed the results and commented on the manuscript.

## Data availability

The evidence data generated and/or analyzed during the current study are available from the corresponding author upon reasonable request.

## Acknowledgments

The authors thank Dr. Hiromu Kashida (Nagoya University) for designing the random oligo DNA sequences, Prof. Akihiro Kusumi (OIST) for advice on single-fluorophore imaging of DPPE, Prof. Hiroaki Tateno for advice on nucleotide conjugation to antibodies, and Dr. Atushi Okazawa (Waseda University) for the support of figure preparation. This research was supported by grants from JST PRESTO JPMJPR15F8, JSPS Grants-in-Aid for Scientific Research (23K08410 to SS; 23K18116 and 23K21299 to HB), and grants from Naito Foundation and Takeda Science Foundation to HB. DeepL, ChatGPT 4o, Grammarly, and Enago, the editing brand of Crimson Interactive Pvt. Ltd supported the English editing of this article.

---

**Supplementary Figure S1.**
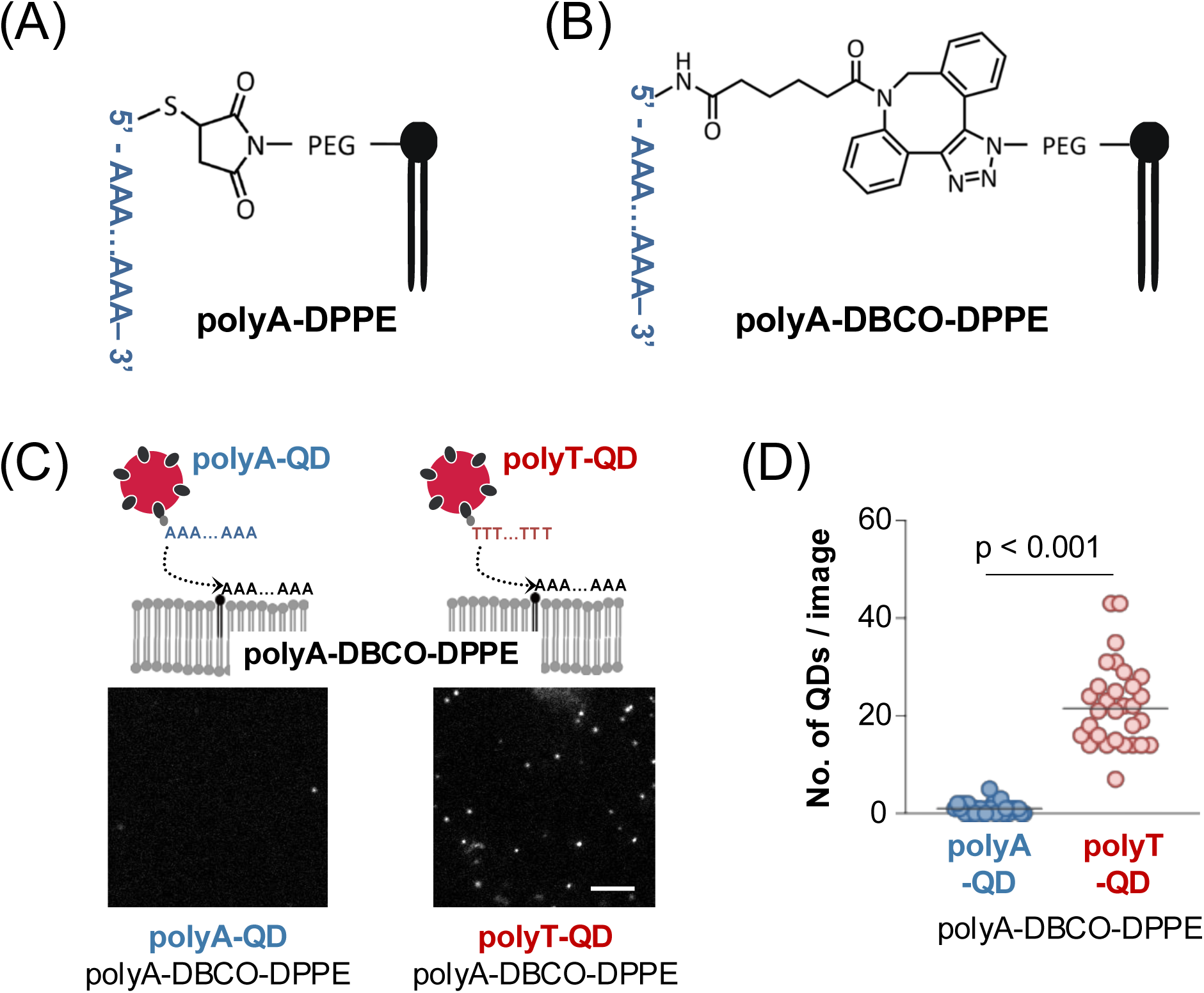
Effect of the linker between DPPE and ssDNA on QD-labeling specificity. (A) PolyA–DPPE generated via thiol–maleimide coupling. (B) PolyA–DPPE–DBCO generated via DBCO–azide click chemistry. (C) Representative images of polyA–DPPE–DBCO labeled with polyA–QD (left) or polyT–QD (right). Bar, 10 µm. (D) Number of QDs per field (41.8 × 41.8 µm; solid lines, medians; n = 29 images for polyA–QD and n = 30 images for polyT–QD). Mann–Whitney U test.

**Supplementary Figure S2.**
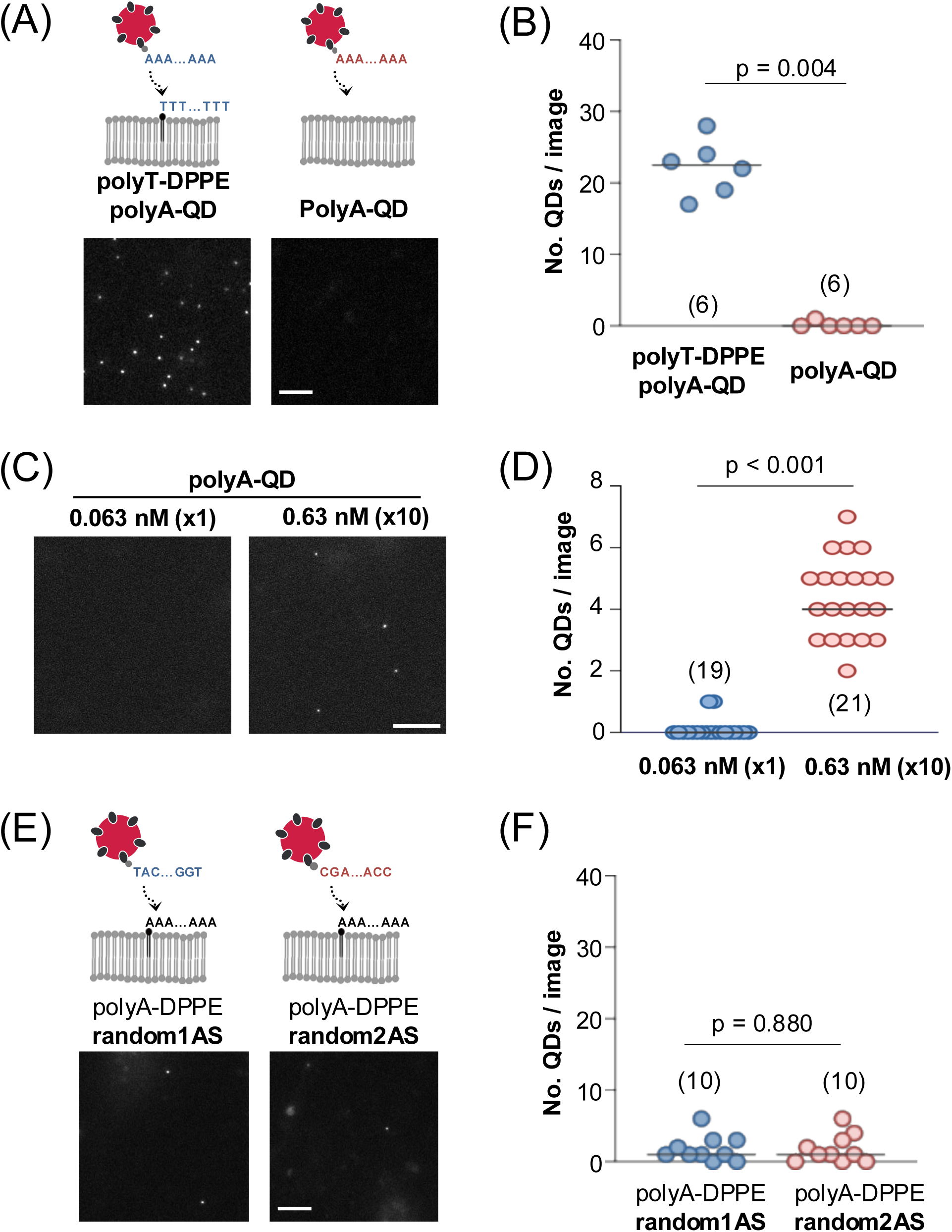
Specificity of ssDNA–QD binding to the DPPE. A) Representative images of neurons incubated with polyA–QD in the presence (left) or absence (right) of polyT–DPPE. (B) Number of QDs per field (41.8 × 41.8 µm; solid lines, medians). (C) Representative images of neurons incubated with 0.063 or 0.63 nM polyA–QD. (D) Number of QDs per field (solid lines, medians). (E) Representative images of neurons expressing polyT–DPPE labeled with QDs carrying random sequences (random1AS or random2AS). (F) Number of QDs per field (solid lines, medians). Bars in (A), (C), and (E), 10 µm. Statistical analysis: (B, D) Mann–Whitney U test. N is shown in parentheses (see Table S2).

**Supplementary Figure S3.**
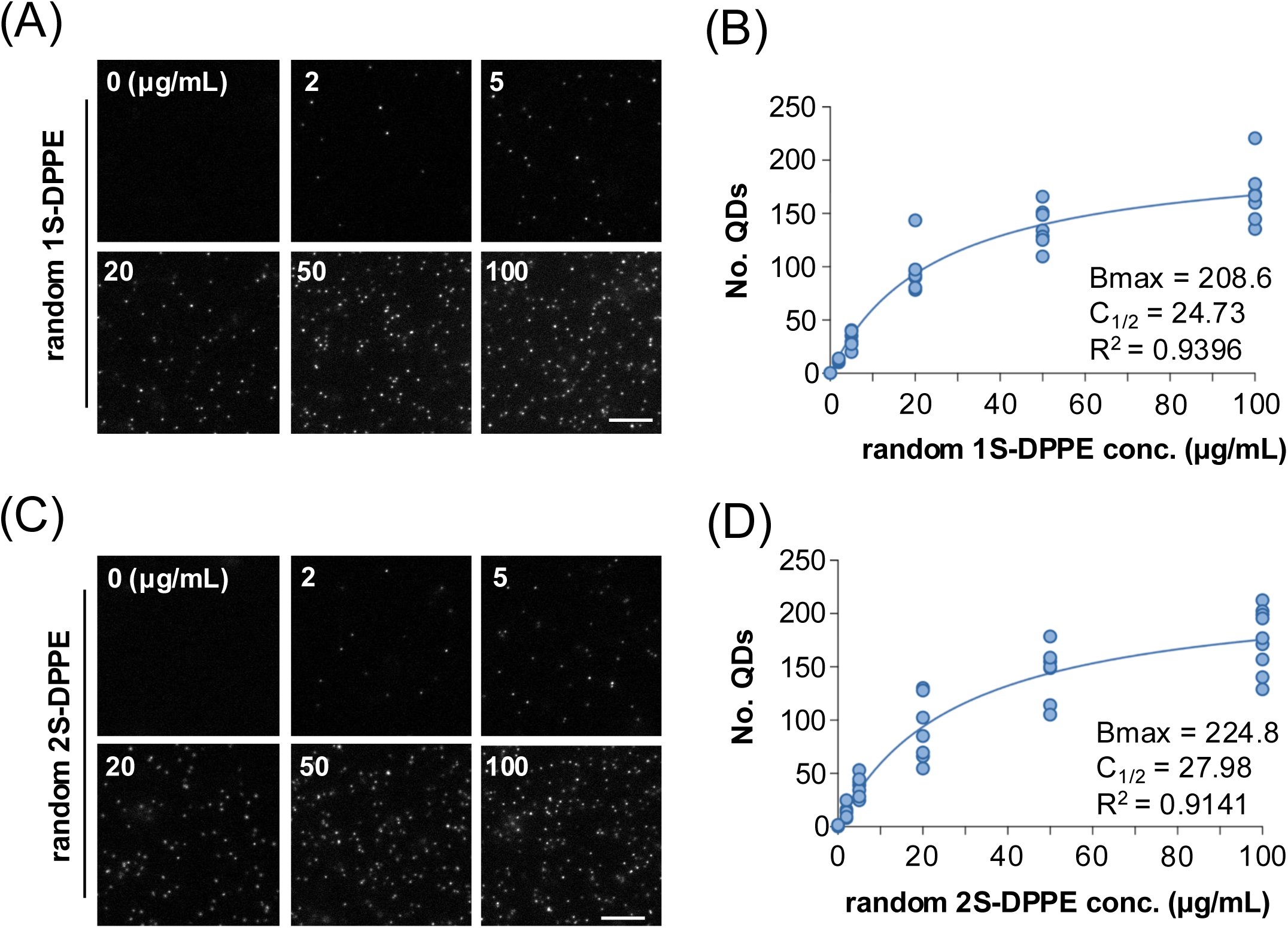
Number of QDs at different concentrations of random 1S-DPPE and random 2S-DPPE. (A, C) Representative QD images for random 1S-DPPE + random 1AS-QD (A) and random 2S-DPPE + random 2AS-QD (C) at the indicated DPPE concentrations (0, 2, 5, 20, 50, 100 μg/mL). Bar, 10 µm. (B, D) Number of QDs per field of view (41.8 × 41.8 µm) for random 1S-DPPE (B) and random 2S-DPPE (D). Each point represents one field. n = 5 images (0 μg/mL) and 7 images (others) for random 1S-DPPE; n = 5 images (0 μg/mL), 7 images (2–50 μg/mL), and 9 images (100 μg/mL) for random 2S-DPPE.

**Supplementary Figure S4.**
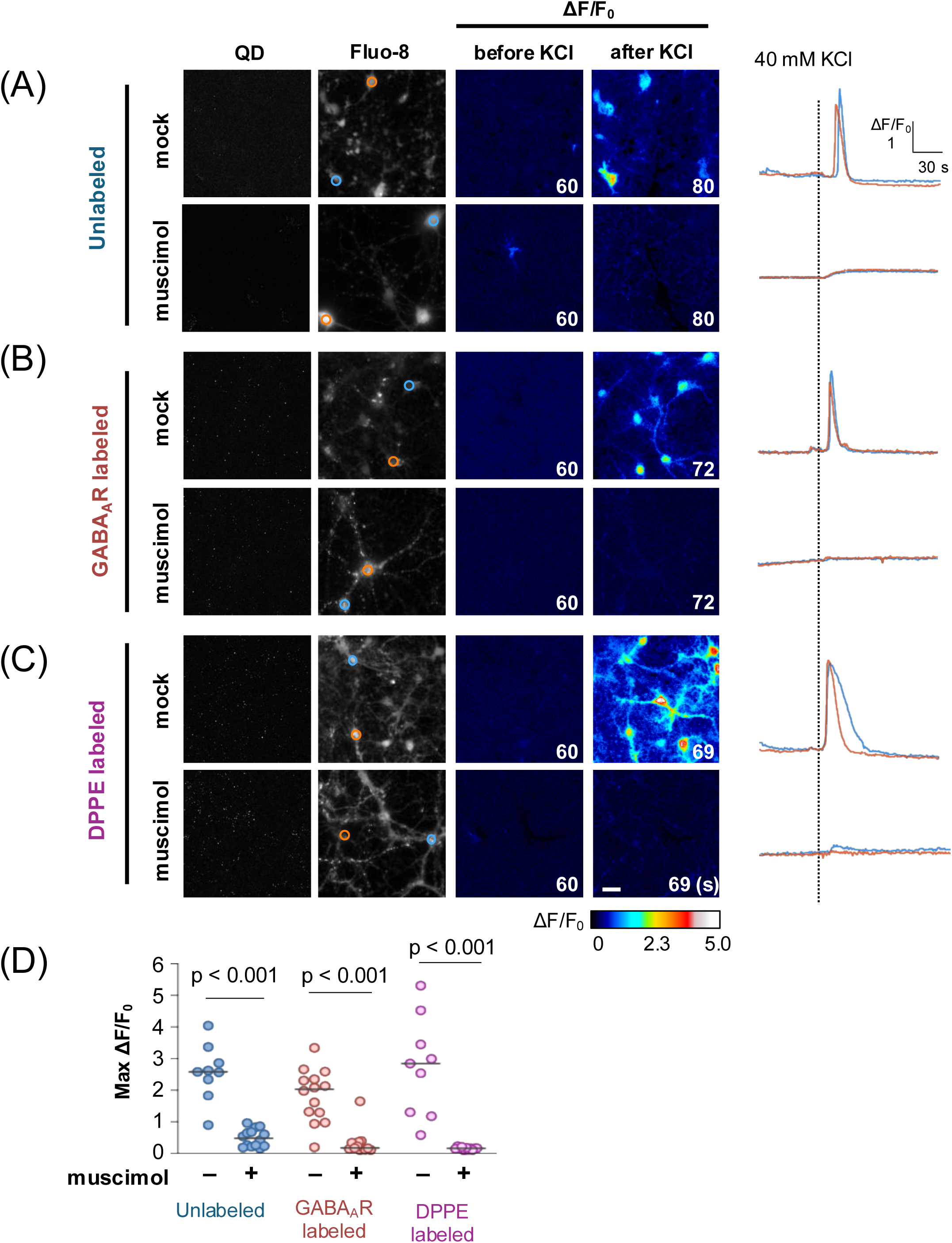
Comparison of KCl-induced intracellular Ca²⁺ responses in the presence or absence of muscimol under QD-labeled and unlabeled conditions. (A–C) QD images, Fluo-8 images, pseudocolor ΔF/F₀ images before and after 40 mM KCl application, and representative ΔF/F₀ traces of neurons with or without preincubation with 20 µM muscimol. Bar, 20 µm. (A) Unlabeled cells; (B) GABA_A_R-labeled cells; (C) DPPE-labeled cells. (D) Maximum ΔF/F₀ with (+) or without (–) muscimol under unlabeled, GABA_A_R-labeled, and DPPE-labeled conditions (solid lines, medians). n = 9 (unlabeled), 14 (unlabeled + muscimol), 14 (GABA_A_R-labeled), 12 (GABA_A_R-labeled + muscimol), 9 (DPPE-labeled), and 11 (DPPE-labeled + muscimol). Statistical analysis: Mann–Whitney U test.

**Supplementary Figure S5.**
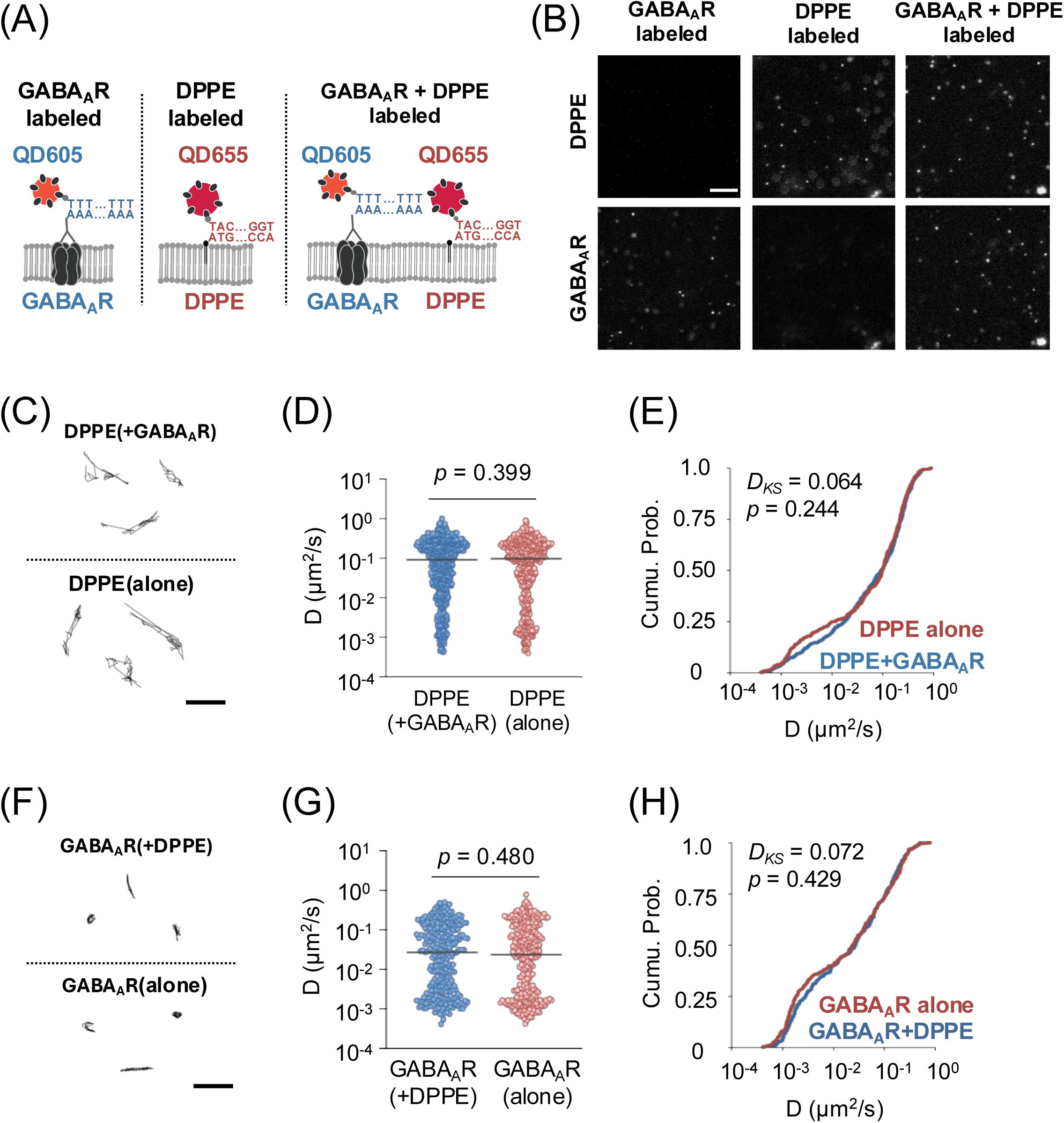
Comparison of diffusion coefficients under single-color and multicolor QD-labeling conditions. (A) Strategy for single-color and multicolor QD labeling of GABA_A_R (QD605) and DPPE (QD655) using oligoDNAs. (B) Representative QD-labeling images. Bars, 10 µm. (C) Representative trajectories of DPPE under single-color and multicolor labeling. Bar, 1 µm. (D, E) Distribution of D for DPPE shown as a dot plot (D) and cumulative distribution (E) (solid lines, medians). n = 451 (DPPE alone) and 611 (DPPE + GABA_A_R). (F) Representative trajectories of QD-GABA_A_R under single-color and multicolor labeling. Bar, 1 µm. (G, H) Distribution of D for GABA_A_R shown as a dot plot (G) and cumulative distribution (H) (solid lines, medians). n = 298 (GABA_A_R alone) and 300 (GABA_A_R + DPPE). Statistical analysis: (D, G) Mann–Whitney U test; (E, H) Kolmogorov–Smirnov test.

**Supplementary Figure S6.**
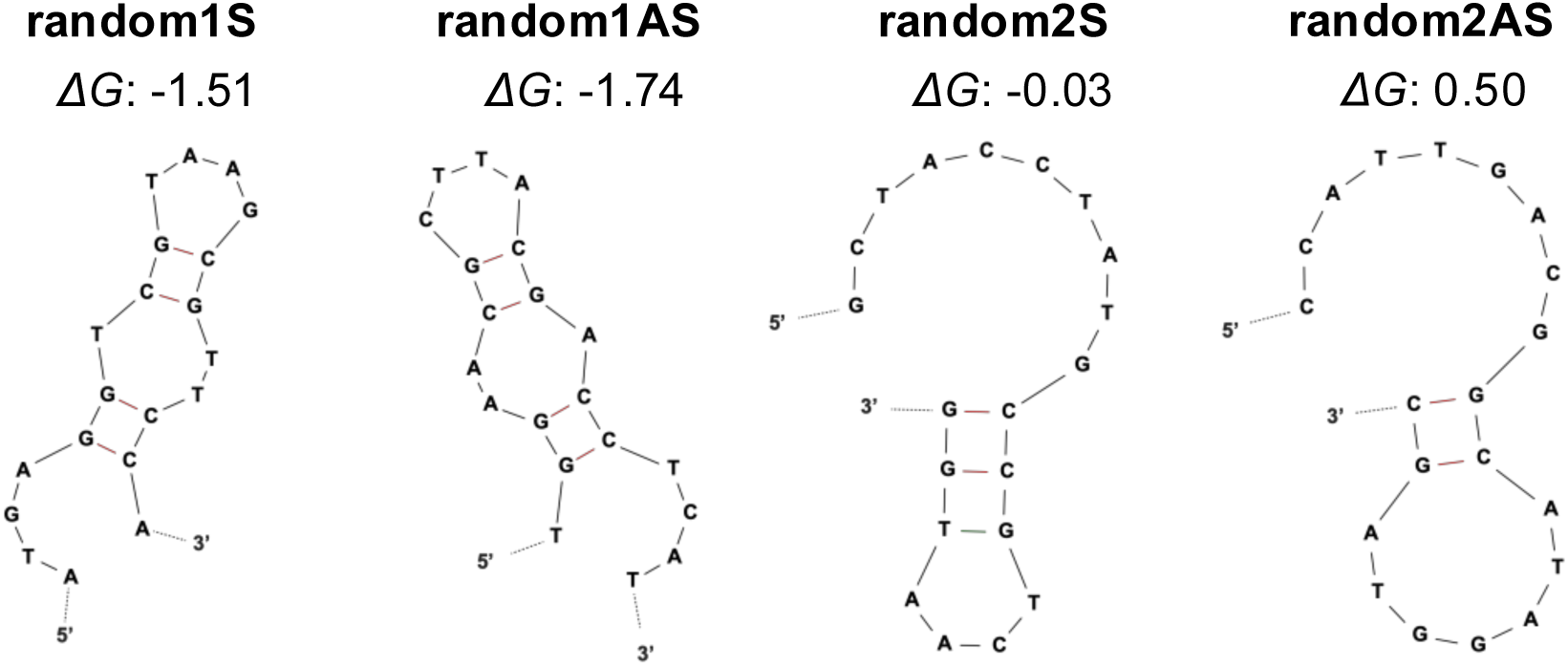
Secondary structures predicted by mfold [33] for the random ssDNAs used in this experiment. Red lines indicate canonical base pairs of Watson–Crick type, which contribute significantly to the thermodynamic stability of the structure. Green lines indicate non-canonical or weak base pairing; ΔG denotes free energy change [kcal/mol].

**Table S1.**
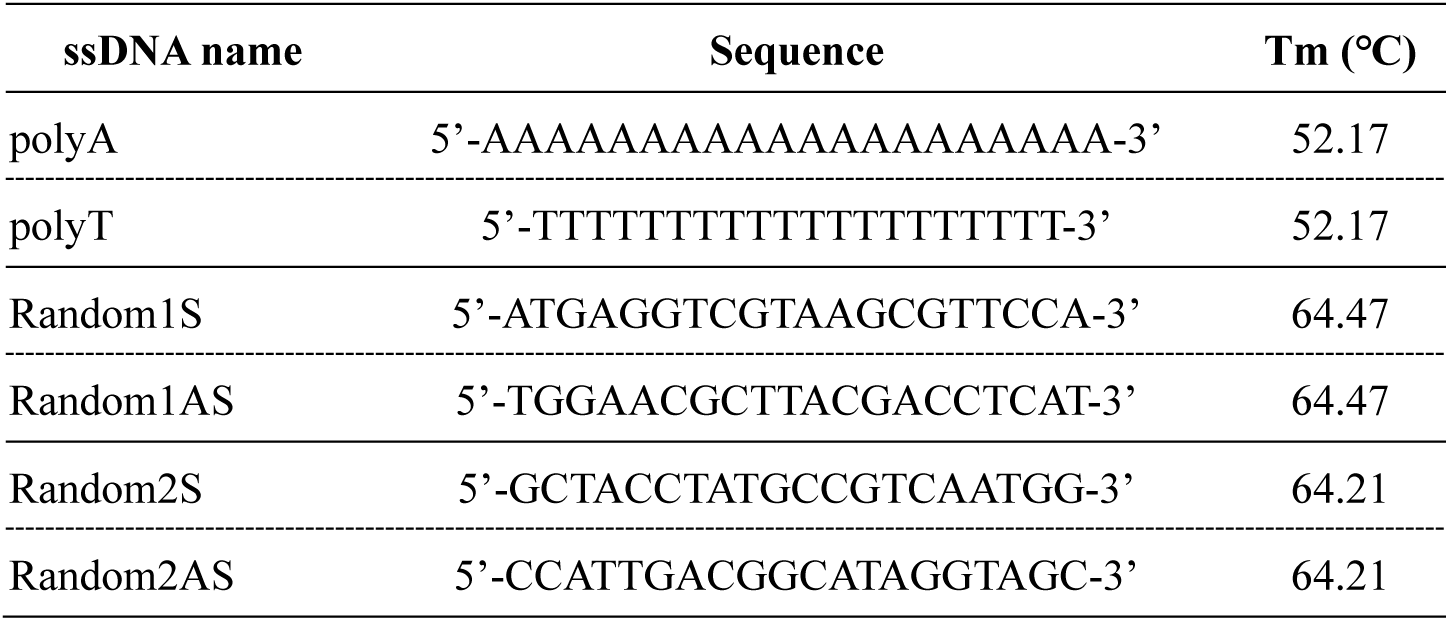
ssDNA nucleotide sequences. List of ssDNA sequences used and melting temperatures (Tm) estimated by nearest neighbor base pairing (Breslauer et al., 1986). PolyT and PolyA, random1S and 1AS, random2S and 2AS, are complementary pairs.

**Table S2:** Summary of the median and Interquartile Range (IQR) for the data. The median and IQR of each data sets are shown here.

| Figure | group | Median<br>(spots/image) | IQR (25% – 75%)<br>(spots/image) | N(images) |
| --- | --- | --- | --- | --- |
| **Figure 2C** | polyA-QD | 1 | 0 – 2 | 10 |
|  | polyT-QD | 241 | 96 – 400 | 10 |
| **Figure 2F** | polyA | 3 | 2 – 4 | 10 |
|  | polyT | 3 | 2 – 4 | 10 |
|  | random1AS | 40 | 26 – 49 | 10 |
|  | random2AS | 1 | 0 – 2 | 10 |
| **Figure 2I** | mock (0min) | 25 | 17 – 29 | 11 |
|  | mock (3min) | 27 | 20 – 30 |  |
|  | DNase (0min) | 31 | 24 – 33 | 11 |
|  | DNase (3min) | 10 | 4 – 11 |  |
| **Figure 4C** | polyA-polyT | 193 | 169 – 206 | 5 |
|  | random1S-1AS | 20 | 17 – 25 | 5 |
| **Figure 5C** | polyA-QD | 2 | 1 – 3 | 30 |
|  | polyT-QD | 28 | 22 – 44 | 30 |
| **Figure S1D** | polyA-QD | 1 | 0 – 1 | 29 |
|  | polyT-QD | 22 | 15 – 26 | 30 |
| **Figure S2B** | polyT-DPPE + polyA-QD | 23 | 20 – 24 | 6 |
|  | polyA-QD | 0 | 0 – 0 | 6 |
| **Figure S2D** | 0.063 nM | 0 | 0 – 0 | 19 |
|  | 0.63 nM | 4 | 3 – 5 | 21 |
| **Figure S2F** | random1AS | 1 | 1 – 3 | 10 |
|  | random2AS | 1 | 0 – 3 | 10 |

| Figure | group | Median<br>($\mu\text{m}^2/\text{s}$) | IQR (25% – 75%)<br>($\mu\text{m}^2/\text{s}$) | N |
| --- | --- | --- | --- | --- |
| **Figure 3C**<br>**3D** | glass attached | $2.40 \times 10^{-2}$ | $1.66 - 3.03 \times 10^{-2}$ | 152 spots |
| | lower | $1.15 \times 10^{-1}$ | $5.70 - 26.6 \times 10^{-2}$ | 368 spots |
| | upper | $2.21 \times 10^{-1}$ | $1.26 - 3.43 \times 10^{-1}$ | 584 spots |
| **Figure 3H**<br>**3I** | lower | $8.23 \times 10^{-2}$ | $0.387 - 20.6 \times 10^{-2}$ | 879 QDs |
| | upper | $1.86 \times 10^{-1}$ | $9.13 - 31.5 \times 10^{-2}$ | 815 QDs |
| **Figure 4E**<br>**4F** | polyA-polyT | $1.38 \times 10^{-1}$ | $7.74 - 22.6 \times 10^{-2}$ | 219 QDs |
| | random1S-1AS | $1.47 \times 10^{-1}$ | $7.86 - 23.0 \times 10^{-2}$ | 253 QDs |
| **Figure 5F**<br>**5G** | ssDNA | $6.49 \times 10^{-2}$ | $1.41 - 12.3 \times 10^{-2}$ | 969 QDs |
| | Fab | $5.19 \times 10^{-2}$ | $0.841 - 12.8 \times 10^{-2}$ | 686 QDs |
| **Figure 6D**<br>**6E** | GABA<sub>A</sub>R | $2.32 \times 10^{-2}$ | $0.300 - 9.56 \times 10^{-2}$ | 419 QDs |
| | DPPE | $1.15 \times 10^{-1}$ | $0.826 - 24.5 \times 10^{-2}$ | 412 QDs |
| **Figure S6D**<br>**S6E** | DPPE (alone) | $9.74 \times 10^{-2}$ | $1.14 - 21.6 \times 10^{-2}$ | 451 QDs |
| | DPPE (+GABA<sub>A</sub>R) | $9.18 \times 10^{-2}$ | $1.78 - 22.9 \times 10^{-2}$ | 611 QDs |
| **Figure S6G**<br>**S6H** | GABA<sub>A</sub>R (alone) | $2.38 \times 10^{-2}$ | $0.19 - 11.1 \times 10^{-2}$ | 298 QDs |
| | GABA<sub>A</sub>R (+DPPE) | $2.72 \times 10^{-2}$ | $0.29 - 10.5 \times 10^{-2}$ | 300 QDs |

| Figure | group | Median (%) | IQR (25% – 75%) (%) | N (images) |
| --- | --- | --- | --- | --- |
| Figure 3G | lower | 17.3 | 4.53 – 25.1 | 28 |
|  | upper | 2.16 | 0 – 7.80 | 28 |

| Figure | group | Median (max $\Delta F/F_0$) | IQR (25% – 75%) (max $\Delta F/F_0$) | N (images) |
| --- | --- | --- | --- | --- |
| Figure S4D | Unlabeled (–muscimol) | 2.58 | 2.34 – 2.86 | 9 |
|  | Unlabeled (+muscimol) | 0.48 | 0.24 – 0.67 | 14 |
|  | GABA<sub>A</sub>R (–muscimol) | 2.03 | 1.30 – 2.34 | 14 |
|  | GABA<sub>A</sub>R (+muscimol) | 0.18 | 0.14 – 0.35 | 12 |
|  | DPPE (–muscimol) | 2.84 | 1.30 – 3.45 | 9 |
|  | DPPE (+muscimol) | 0.16 | 0.13 – 0.18 | 11 |

